# Learning generalizable visuomotor mappings fo*r de novo* skills

**DOI:** 10.1101/2023.07.18.549179

**Authors:** Carlos A. Velázquez-Vargas, Nathaniel D. Daw, Jordan A. Taylor

## Abstract

A fundamental feature of the human brain is its capacity to learn novel motor skills. This capacity requires the formation of vastly different visuomotor mappings. In this work, we ask how these associations are formed *de novo*, hypothesizing that under specific training regimes generalizable mappings are more readily formed, while in others, local state-actions associations are favored. To test this, we studied learning in a simple navigation task where participants attempted to move a cursor between various start-target locations by pressing three keyboard keys. Importantly, the mapping between the keys and the direction of cursor movement was unknown to the participants. Experiments 1 and 2 show that participants who were trained to move between multiple start-target pairs had significantly greater generalization than participants trained to move between a single pair. Whereas Experiment 1 found significant generalization when start-targets were distal, Experiment 2 found similar generalization for proximal targets, which suggests that generalization differences are due to knowledge of the visuomotor mapping itself and not simply due to planning. To gain insight into the potential computational mechanisms underlying this capacity, we explored how a visuomotor mapping could be formed through a set of models that afforded construction of a generalizable mappings (model-based), local state-action associations (model-free), or a hybrid of both. Our modeling work suggested that without continued variability between start-target pairs during training, model-based processes eventually gave way to model-free processes. In Experiment 3, we sought to further test this shift in learning processes by exposing participants to initially high variability before settling into a condition of no variability over a long-period of training. We found that generalization performance remained intact after a prolonged period of no variability suggesting that the formation of visuomotor mappings might occur at an early stage of learning. Finally, in Experiment 4 we show that adding stochasticity in the mapping can also promote model-based learning of a visuomotor mapping, suggesting that the learning may unfold implicitly. Overall, these studies shed light on how humans could acquire visuomotor mappings in their lives through exposure to variability in their feedback.

## Introduction

The first problem to be overcome in learning any novel motor skill is to associate particular actions with desired outcomes. This problem has become increasingly complex in the digital age, where the mapping between actions and outcomes can be as diverse as the imagination allows – just consider the variety of action-outcome associations underlying digital applications and video games. For example, using two thumbs to type a text message, using a pinch motion to zoom in and out of content on a smartphone, or steering a car in a video game. At first, learning these novel mappings is cumbersome and effortful but as learning progresses a mapping between actions and outcomes is eventually formed, allowing the individual to use the device successfully with ease. The formation of this mapping is arguably one of the most important steps for learning any new motor skill (Fitts and Posner 1967; Adams, 1971; Ackerman, 1988; Newell, 1985, 1991). Surprisingly, however, we know very little, with a few exceptions (Mosier et al., 2005; Liu et al., 2011), about how novel motor mappings are established *de novo*.

Traditionally, the question of how motor mappings are learned has been the focus of sensorimotor adaptation tasks (e.g., prisms, visuomotor rotations, and force fields), which impose a perturbation on the sensory outcome of a movement (Shadmehr and Mussa-Ivaldi, 1994; Martin et al., 1996; Krakauer et al., 2000). While adaptation tasks were originally thought to serve as a model paradigm to study this question (Jordan and Rumelhart, 1992; Miall and Wolpert, 1996; Shadmehr et al., 2010), in recent years, it has become clear that these tasks may only pressure the recalibration of an existing motor mapping when faced with an externally imposed perturbation – not the establishment of the mapping in the first place (Krakauer et al., 2019; Hadjiosif et al., 2021). Only when these recalibration mechanisms fail to fully counteract these perturbations in adaptation paradigms (Hadjiosif et al., 2021), may *de novo* learning engage to develop a new controller for the task (Yang et al., 2021).

While there have been numerous studies of operant conditioning and associative learning, linking actions to outcomes, it is unclear the degree to which learning in these studies reflects the formation of a motor mapping *per se* (Thorndike, 1927; Petrides, 1997; Elsner and Hommel, 2004). It can be helpful, at least conceptually, to distinguish two levels of choice: a more abstract, internal level of reasoning about goals and state changes, and a more external, response-focused level about how to use movements to bring these plans to fruition. In many studies, such as in spatial or maze navigation, the agent already knows the control policy of how to move (i.e., how an action leads to a state change) and instead the focus is on reasoning or learning at a more abstract level how the state change leads to a desired outcome in terms of reward (Sutton and Barto, 1998; Simon and Daw, 2011). Conversely a different set of paradigms focus entirely on externally-cued responses, without any internal plan. Such tasks include motor sequence learning (Nissen and Bullemer, 1987; Willingham et al., 1989; Curran and Keele, 1993), discrete sequence production (Verwey, 2001; Abrahmase et al., 2013), and *m x n* tasks (Hikosaka et al., 1995; Bapi et al., 2000), all of which can be viewed as a form of *de novo* motor learning, establishing a relationship between arbitrary actions and outcomes. However in these studies, there is no underlying mapping from internal goals, from which a generalizable, motor map may form. Generalization to new situations or contexts is considered a hallmark feature of a motor mapping, as opposed to rote memorization of stimulus-response associations (Shadmehr and Mussa-Ivaldi, 1994; Mussa-Ivaldi, 1999; Shadmehr, 2004).

Furthermore, the different variants of these sequence learning tasks are externally generated, such that the appropriate sequence of responses is fully specified by the experimental stimuli (Bera et al., 2021). The participant must precisely follow the set of stimulus-response pairs to be successful in the task. As such, they may only reflect a subset of the kinds of motor skills that we perform in everyday life that are internally generated. Thus, while there has been tremendous progress in understanding how externally-generated, stimulus-response mappings are learned, there has been comparatively less progress in understanding how internally-generated, response-outcome mappings are formed.

The increased complexity and degrees of freedom available when learning an internally-generated, response-mapping may be one potential reason why progress has been slow. The difficulty is in designing and studying a task that is in the “Goldilocks zone” between sufficient experimental complexity and analytical tractability (Opheusden and Ma, 2019). Fermin and colleagues developed the *grid-sailing task* to fit within this zone to study the core problem of learning an internally-generated mapping de novo: The formation of a novel and arbitrary motor mapping (Fermin et al., 2010, 2016). Here, participants learn to navigate a cursor from various starting locations to various target locations on a grid through a series of keypresses. The goal is to navigate to the target location in the minimum number of moves as quickly as possible. Importantly, there is an unintuitive and arbitrary mapping between the keys and cursor movement that must be learned (i.e., the control policy).

While participants can learn this task within a relatively short period, it remains an open question as to how this is accomplished (Fermin et al., 2010; Bera et al., 2021; Dundon, 2021). The formation of a new mapping is not always guaranteed. If the task only demands the repetition of a limited set of actions, then only local state-action associations may be learned – a form of rote memorization, which is likely what occurs in studies of sequence learning. However, if there is a greater degree of variability in training, then a richer representation of skill may be learned, such as the formation of an internal model between the action-outcome space. This would afford the ability to generalize outside the range of training (Schmidt 1975; Newell and Shaprio, 1976; McCracken and Stelmach, 1977; Kerr and Booth, 1978; Catalano and Kleiner, 1984; Berniker et al., 2014) – an idea that echoes classic theories of stimulus variability in learning (Estes and Burke, 1953; Raviv et al., 2022). These two forms of learning mirror the instance-based and algorithmic processes of a classic theory for automatization (Logan 1988), as well as the more modern notions of model-free and model-based reinforcement learning (Sutton and Barto, 1998; Daw et al., 2005, 2011; Haith and Krakauer, 2013; Raviv et al., 2023). In particular, the latter formalism seems well suited to capturing the candidate mechanisms. Model-based reinforcement learning is well suited to capture the covert formation of an abstract, internal plan that can then be generalizably realized through a separately learned motor mapping. This leads to the hypothesis that, much as in other circumstances such as operant leverpressing (Daw et al., 2005), simpler model-free (stimulus-response) learning will instead dominate when a narrow range of actions is overtrained.

Here, through a series of experiments, we seek to build on this work by leveraging the grid-sailing task as a model paradigm to study how novel and arbitrary motor mappings are learned *de novo* and characterizing how they may be learned through the model-free and model-based reinforcement learning framework. We hypothesize that the formation and representation of a novel motor mapping depends on the particular conditions of training. Specifically, the degree of exploration between the number of potential action goals and possible solutions to achieve that goal may pressure formation of a generalizable motor mapping over local state-action associations (e.g., rote memorization of specific sequences of actions). Generalization to untrained conditions will provide a key test for the existence of a motor mapping.

## Results

### Experiment 1

#### Behavioral Results

How does training variability constrain learning and generalization of a visuomotor mapping? Two groups of participants (n=32) performed a grid navigation task (Figure 1A) where they moved a cursor from various start to target locations using the J, K and L keys of a standard keyboard (see *Methods* for details). In the Single group, participants trained to move between a single start-target pair, while in the Multiple group participants trained to move between multiple start-target pairs (Figure 1B). We predicted that performance improvements for the Multiple group would be slower during training, but they would be able to generalize their performance to novel start-target pairs, reflecting the formation of a key-to-direction mapping rather than local state-action associations. In contrast, the Single group would show faster performance improvements during training, but be unable to generalize to new start-target pairs. Participants performed 260 training trials followed by 20 trials of generalization interleaved with 20 training trials.

**Figure 1:**
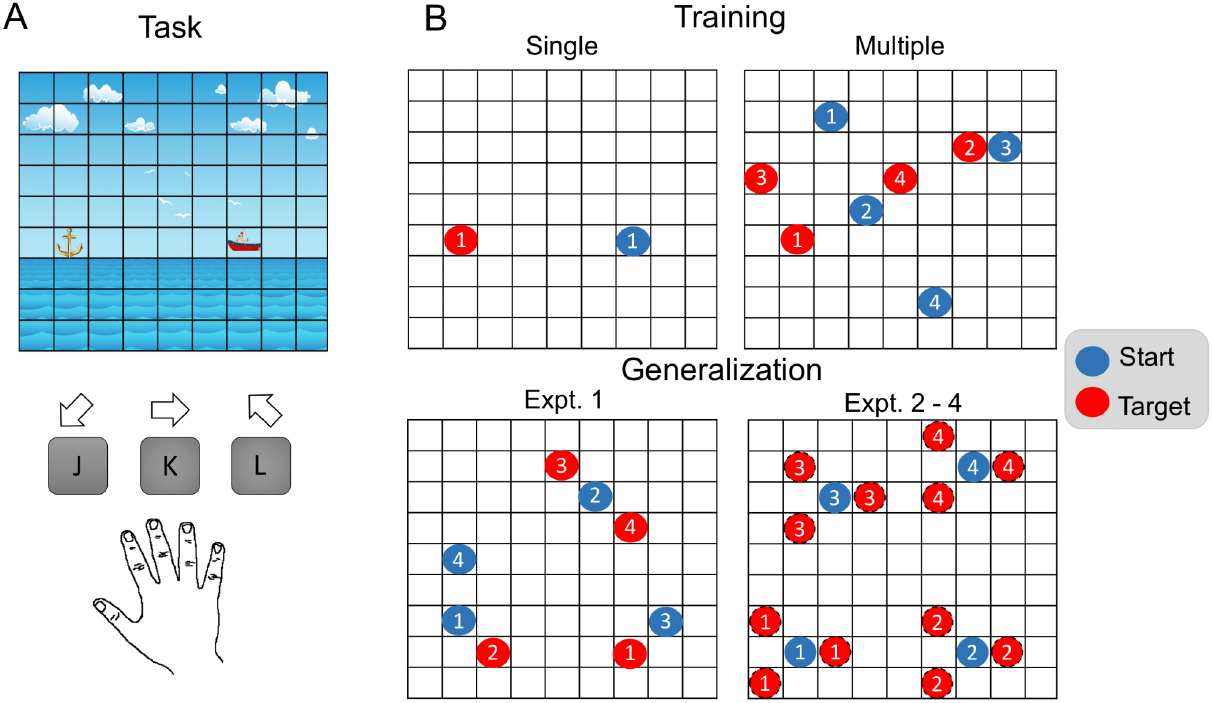
Experimental task. **A)** Participants moved a cursor (ship) from start to target (anchor) locations in a grid environment. For Experiments 1-3, participants used a deterministic action mapping of three keys with moving directions: bottom-left, right and top-left. In Experiment 4, the key-to-direction mapping randomly changed after each keypress with a probability of 0.2 to any of the remaining directions. **B)** In the four experiments, participants were trained with a single or multiple start-target pairs (see Methods for details). A generalization phase was presented after training where the target locations were either seven (Experiment 1) or one move away (Experiments 2-4) from the starting point. Blue and red circles represent the start and target locations and the specific start-target pairs are indicated with numbers. Only one pair was presented per trial.

Figure 2A shows the proportion of optimal arrivals over trials for both groups, where an optimal arrival is a trial where subjects arrive at the target using the minimum number of key presses (7 moves). As expected, the Multiple group had a slower learning curve, however, both the Single and Multiple groups reached the same level of performance by the end of the training phase (comparison of the last bin of training trials between groups; t(29.06) = -0.5; p = 0.61; Figure 2B). Of more importance is how well the groups perform when new start-target pairs are introduced in the generalization phase (Figure 2C). Here, we performed a two-step analysis where we first compared performance in the generalization phase to performance at the end of the training phase before comparing potential differences in generalization between the groups. For the Single group, we found that performance was significantly worse at the onset of the generalization phase compared to the end of the training phase (t(15) = -1.21; p < 0.001). In contrast, the Multiple group’s performance during generalization was similar to the end of the training phase (t(15) = -1.21; p = 0.242). When comparing between the groups, it was clear that the Multiple group performed significantly better than the Single group (t(29.9) = -5.1; p < 0.001; Figure 2D), even when controlling for multiple comparisons (t(29.34) = -3.5; p = 0.001; Figure 2E).

**Figure 2:**
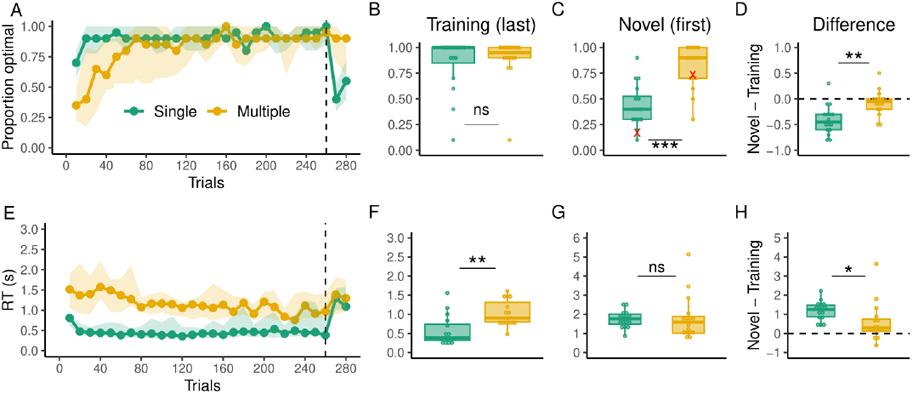
Behavioral results of Experiment 1. **A)** Proportion of optimal arrivals over trials for the Single (green) and Multiple (gold) groups. The dashed line indicates the beginning of the generalization phase. The solid dotted line represents the median and the shading the interquartile range. **B)** Proportion of optimal arrivals in the last bin of the training phase. **C)** Proportion of optimal arrivals in the first bin of generalization (novel pairs). Red marks indicate performance in the very first trial of generalization for all subjects. **D)** Difference in the proportion of optimal arrivals between the first bin of generalization and the last one of training. The dashed line here indicates no performance change from training to generalization **E)** RTs over trials. **F)** RTs in the last bin of training trials. **G)** RTs in the first bin of generalization **H**) Difference in RTs between the first bin of generalization and the last one of training.

We also examined how planning (reaction time, RT) evolved over training. RTs were overall higher in the Multiple group group throughout training (Figure 2D; t(29.99) = -4.82; p < 0.001) and higher at the last bin of training trials (figure 2F; t(21.21) = -2.85; p = 0.009) but not during the first bin of generalization trials (Figure 2G; t(19.36) = -0.17; p = 0.86). There was an increase in RTs from the last training bin to the first generalization bin in the Single (t(15) = 9.54; p < 0.001) and Multiple groups (t(15) = 2.34; p = 0.03), however, this increase was significantly greater in the Single group (Figure 2D; t(21.96) = 2.14; p = 0.043). In addition, the groups only differ in RTs over training, but not in any of the inter-key-intervals (Figure S1). Overall, greater generalization in the Multiple group together with higher RTs suggest that this group could have relied more heavily on model-based planning computations involving the visuomotor mapping of the task as opposed to the Single group which could have developed less computationally demanding state-action associations.

#### Modeling Results

In order to explore the cognitive processes that give rise to the differences in performance between the groups, we evaluated three different computational models. On one end of the spectrum, we tested a model-free reinforcement learning algorithm which learns state-action values on the grid using prediction errors. On the other end of the spectrum, we implemented a model-based algorithm that learns the visuomotor mapping of the task and uses it to find the shortest route to the target using tree search. We selected these models as they make contrasting predictions about generalization performance in the task. A model-free reinforcement learning algorithm would generate chance-level responses for states that have not been experienced in the past, thus predicting poor generalization. A model-based algorithm would instead be able to generalize once the visuomotor mapping has been learned. Finally, we tested a hybrid between the model-free and model-based algorithms, which is a weighted sum of the two components on each trial.The relative weighting is given by a timeseries of free parameters, and can evolve over time. We used the logarithm of the marginal likelihood (LML) as a metric for model comparison (Table 1).

**Table 1:** Model comparison using the Logarithm of the Marginal Likelihood (LML) for Experiment 1. 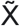 indicates the median value across subjects, and inside the square brackets we show the interquartile range. In the third column we show the number of subjects that according to LML were best described by a given model.

| Groups | Models | LML | LML<br>wins |
| --- | --- | --- | --- |
| Single | MF | $\tilde{X} = -1560[-1919,-1397]$ | 8/16 |
| | MB | $\tilde{X} = -1985[-2279,-1672]$ | 0/16 |
| | Hybrid | $\tilde{X} = -1581[-2087,-1404]$ | 8/16 |
| Multiple | MF | $\tilde{X} = -1768[-2110,-1529]$ | 0/16 |
| | MB | $\tilde{X} = -1556[-2027,-1317]$ | 5/16 |
| | Hybrid | $\tilde{X} = -1539[-1964,-1300]$ | 11/16 |

For the Single group, half of the participants were best explained by the model-free algorithm, whereas the other half were best explained by the Hybrid model. In contrast, for the Multiple group, 31% (5 out of 16) of participants were best explained by the model-based algorithm whereas 69% (11 out of 16) were best explained by the hybrid model. Using the LML values, we also obtained model comparison metrics at the group level (Stephens, Friston, 2009, 2016, see methods for details; Figure 3A). According to this analysis, a random subject taken from the Single group would have a 51% probability of being best described by the model-free algorithm and 49% of being best described by the hybrid model. For the Multiple group, there is a 26% and 74% probability that a random subject is best described by the model-based and hybrid algorithms, respectively.

**Figure 3:**
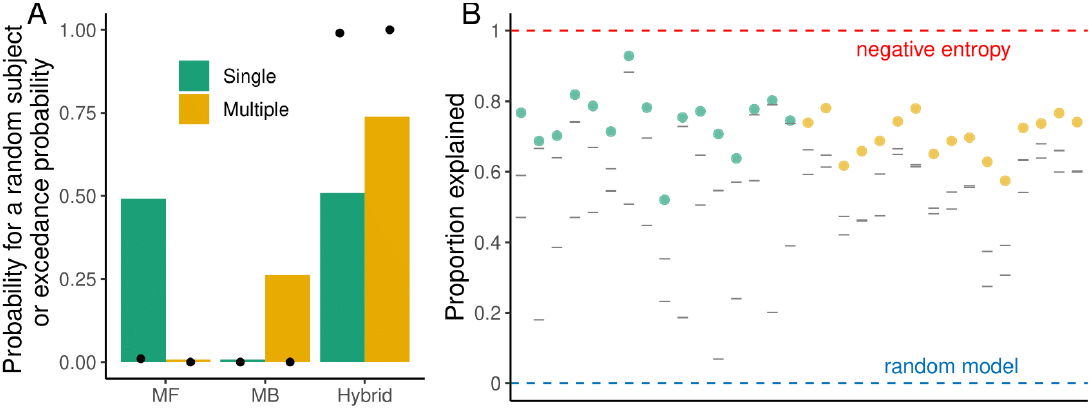
Group model comparison and absolute goodness of fit for Experiment 1. **A:** Probability that a random subject taken from the Single and Multiple groups is best described by the tested models. Black dots indicate the exceedance probability. **B:** Proportion of the variability in the data explained by the models. Colored dots represent this value for the model with the best estimate of the negative cross entropy (see Methods for details) and gray lines represent the values for the other two models. Red and blue dashed lines represent the negative entropy (upper boundary) and the performance of a random model (lower boundary).

We also computed the probability that a given model is more likely than the others in the population, i.e. the exceedance probability. According to this metric, there is a high probability (>99%) that a hybrid model is better than the model-based and model-free in each group. In addition, in order to evaluate how good the models were in the absolute sense, we computed the proportion of the variability in our data that was explained by our models as compared to the negative entropy, which is the upper boundary for any probabilistic model (see Methods for details). The median proportion of the variability explained by the best model in the Single group was 74%, whereas in the Multiple group it was 70% (Figure 3b).

Given that the hybrid best explained the majority of the participants’ data in both the Single and Multiple groups, we analyzed the estimated weights toward the model-based component across trials to see if there was any difference between the groups. Starting from trial 60 and until the end of the training phase, the weight toward the model-based component in the Multiple group was significantly higher (Figure 4). Similarly, the model-based weight was significantly higher in the Multiple group in the first ten trials of generalization compared to the Single group. However, the Single group appeared to switch to a predominantly model-based mode of operation, which would be the only way in this model for performance to improve quickly when encountering new start-target pairs and is also consistent with the change in reaction times. *De novo* model-based learning would have been possible since the target location was 7 moves away from the starting location and participants were provided move-by-move feedback about the consequences of each button press. For this reason, it is unclear if participants in the Single group were using feedback to learn the mapping during the generalization phase or if during the generalization phase they reverted to a memory of the mapping already learned covertly during training, which we address in Experiments 2 and 3, respectively.

**Figure 4:**
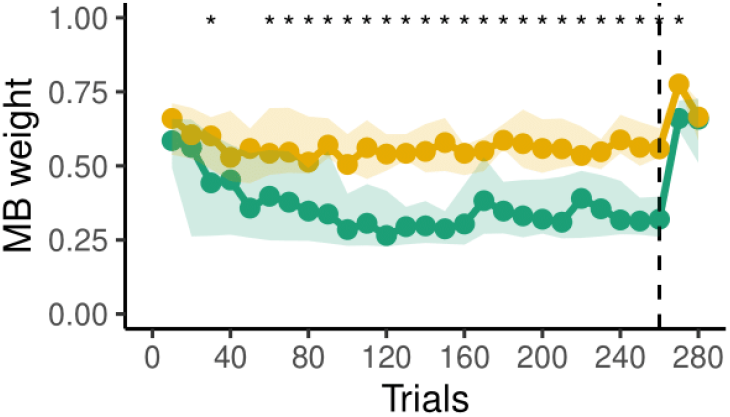
Model-based weighting of the hybrid model over trials for the Multiple (gold) and Single (green) groups of Experiment 1. A value of 1 reflects fully model-based, while a value of 0 reflects fully model-free. The dashed line demarcates the start of the generalization phase.

### Experiment 2

In Experiment 1, we found that the Multiple group readily generalized their performance to new start-target pairs during the generalization phase, while the Single group struggled to generalize early on during generalization. However, feedback was available during the generalization phase, allowing the Single group to learn the mapping to recover performance, which is suggested by their elevated RTs during the generalization phase. Here, we sought to test this possibility by not providing feedback during the generalization phase and placing the target only one move away from the starting location (Figure 1B). In addition, to prevent learning during the generalization phase we removed the interleaved training trials. By removing the sequential, multiple responses to arrive at the target in the generalization phase, we could also rule out a potential confound due to planning of the sequential movements. If participants in the Single group still underperform the Multiple group in this simple situation, it would provide further evidence that they did not know or could not use the mapping to the extent that the Multiple group could, by the end of the training phase.

#### Behavioral Results

Similar to Experiment 1, our analysis focuses on optimal arrivals and RT. Figure 5A shows that the Multiple group has a slower learning curve but both groups reached the same level of performance by the end of the training phase (Figure 5B; t(29.99) = -0.19; p = 0.84). Of primary interest is how each group performed during the generalization phase where only 1 move was required and no feedback was provided. We first compared performance in the generalization phase with performance at the end of training for each group separately before looking for differences in generalization between the groups. The Single group’s performance was significantly worse at the onset of generalization (first bin; t(15) = -5.03; p < 0.001) compared to late in training. On the other hand, the Multiple group’s performance did not significantly decrease between the training and generalization phases (t(15) = -1.24; p = 0.23), and outperformed the Single group early (first bin; t(22.68) = -4.81; p < 0.0001) and late in the generalization phase (second bin; t(19.41) = -3.1; p = 0.005). Notably, the performance of the Single group still remained greater than chance (XX statistical test), which suggests that even though they performed worse than the Multiple group, they could have recalled some knowledge about the mapping. The continued worse performance by the Single group in the generalization phase suggests that they likely did not develop or maintain the mapping and suggests that variability in training may be necessary for a model-based algorithm to be trained.

**Figure 5:**
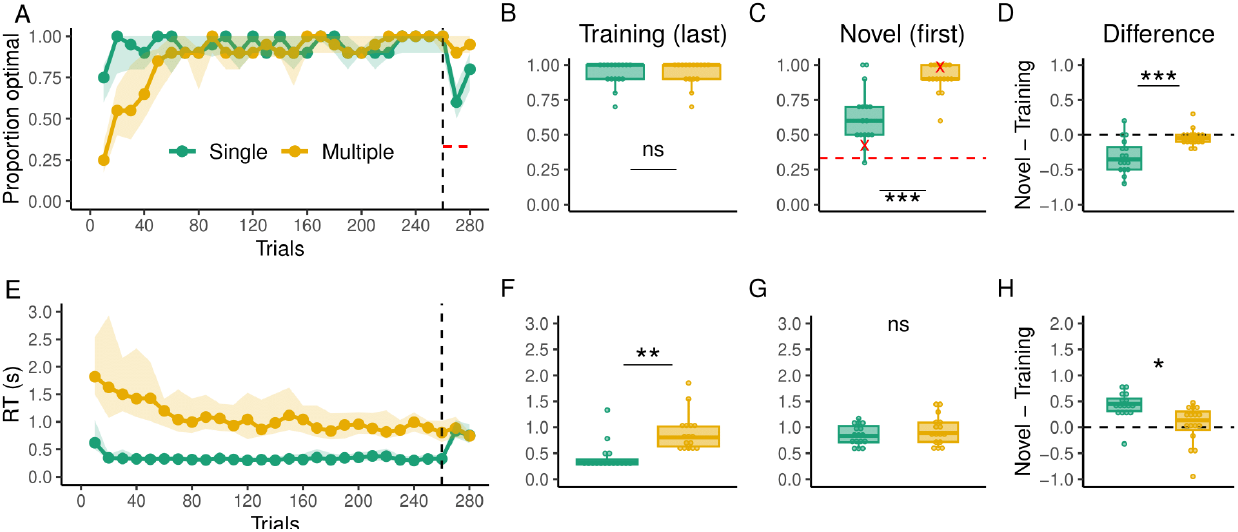
Behavioral results of Experiment 2. **A:** Proportion of optimal arrivals over trials for the Single (green) and Multiple (gold) groups. The black dashed line indicates the beginning of generalization trials. The red dashed line indicates chance level of performance. The solid dotted line represents the median and the shading the interquartile range. **B:** Proportion of optimal arrivals in the last bin of training trials. **C:** Proportion of optimal arrivals in the first bin of generalization (novel pairs). Red marks indicate performance in the very first trial of generalization for all subjects. **D:** Difference in the proportion of optimal arrivals between the first bin of generalization and the last one of training. The dashed line here indicates no performance change from training to generalization **E:** RTs over trials. **F:** RTs in the last bin of training trials. **G:** RTs in the first bin of generalization **H**: Difference in RTs between the first bin of generalization and the last one of training.

Similar to Experiment 1, RTs in the Multiple group were overall higher during training (Figure 5E; t(16.06) = -8.31; p < 0.001) and higher at the last bin of training trials (Figure 5F; t(28.13) = -4.22; p < 0.001) but not in the first bin of generalization trials (Figure 5G; t(26.48) = -1.08; p = 0.28). RTs significantly increased from the last training bin to the first generalization bin in the Single (t(15) = 6.55; p < 0.001) but not in the Multiple group (t(15) = 0.41; p = 0.68). In addition, this change in RTs was significantly greater in the Single group (t(26.43) = 3.34; p = 0.002). The differences in RTs increase suggest that the Multiple group could readily use the mapping to arrive at the novel locations without the need to deploy further computational resources, whereas the Single group had to switch from state action associations to learn the mapping.

#### Modeling Results

As in Experiment 1, we evaluated a model-free, a model-based and a hybrid algorithm. The results of the model comparison are shown in Table 2. According to the LML, in the Single group, 69% (11 out of 16) of the participants were best explained by the model-free algorithm, 19 % (3 out of 16) by the Hybrid model and 12% (2 out of 16) by the model-based algorithm. In the Multiple group, 56% (9 out of 16) of our participants were best explained by the model-based algorithm, 37 % (6 out of 16) by the Hybrid model and 7 % (1 out of 16) by the model-free algorithm.

**Table 2:** Model comparison using the Logarithm of the Marginal Likelihood (LML) for Experiment 2. 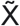 indicates the median value across subjects, and inside the square brackets we show the interquartile range. In the third column we show the number of subjects that according to LML were best described by a given model.

| Group | Models | LML | LML<br>wins |
| --- | --- | --- | --- |
| Single | MF | $\tilde{X} = -1233[-1513,-967]$ | 11/16 |
| | MB | $\tilde{X} = -1604[-1743,-1282]$ | 2/16 |
| | Hybrid | $\tilde{X} = -1321[-1528,-1097]$ | 3/16 |
| Multiple | MF | $\tilde{X} = -1355[-1632, -1164]$ | 1/16 |
| | MB | $\tilde{X} = -1220[-1392, -1083]$ | 9/16 |
| | Hybrid | $\tilde{X} = -1220[-1446, -1057]$ | 6/16 |

In addition, as part of our model comparison at the group level, in Figure 6A we show that a random subject taken from the Single group would have a 74% probability of being best described by the model-free algorithm, 19% of being best described by the hybrid model and 7% by the model-based algorithm. In contrast, in the Multiple group, there is a 59%, 35% and 6% probability that a random subject is best described by the model-based, hybrid and model-free algorithms, respectively. According to the exceedance probability, there is a high probability (>99%) that a model-free algorithm is better than the other models in the Single group, whereas in the Multiple group there is a high probability (>99%) that the model-based algorithm is better than the other models. Thus, while the predominant models were categorical MB and MF in the current experiment (and the hybrid model less often favored, compared to the previous experiment), importantly, some participants were still best described by a Hybrid model in both the Single and Multiple groups. In Figure S2 we show the dynamics of the MB component over trials (which even for subjects best fit by MB or MF is useful descriptively for examining the dynamics of the model evidence). This has a similar trend as in Experiment 1, being higher for the Multiple group during training and early in generalization. This corroborates our hypothesis that the mapping was better learned or used by participants in the Multiple group.

**Figure 6:**
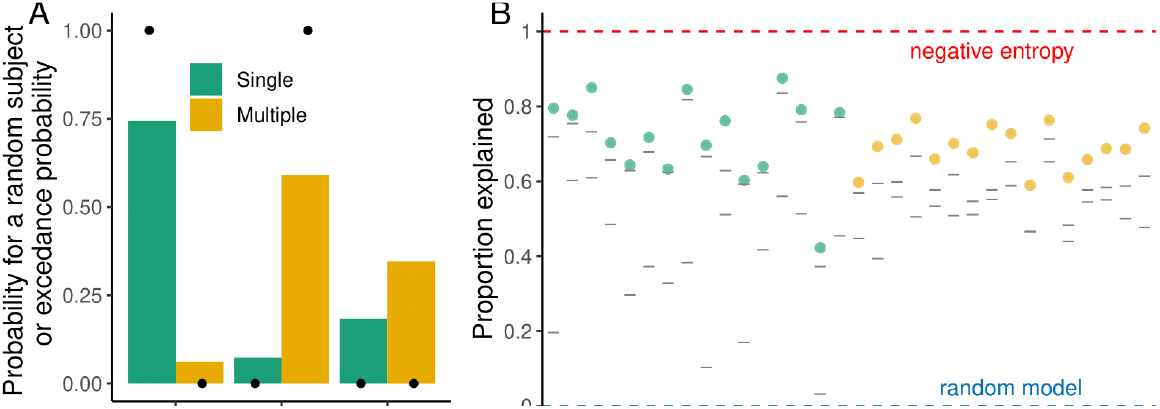
Group model comparison and absolute goodness of fit for Experiment 2. **A:** Probability that a random subject taken from the Single and Multiple groups is best described by the tested models. Black dots indicate the exceedance probability. **B:** Proportion of the variability in the data explained by the models. Coloured dots represent this value for the model with the best estimate of the negative cross entropy and gray illness represent the values for the other two models. Red and blue dashed lines represent the negative entropy (upper boundary) and the performance of a random model (lower boundary).

### Experiment 3

The behavioral and modeling results from Experiments 1 and 2 when taken together, suggest that training variability, as in the Multiple group, strongly encourages learning of the visuomotor mapping rather than just local state-action associations (e.g., simple rote memorization of a sequence of actions), as in the Single group. The modeling results also revealed that there was a tendency for the model-free algorithm to gain preference for behavioral output as training continued, which could reflect an economical choice to limit the computational demands of using a model-based algorithm. However, in Experiment 1, when the Single group was exposed to the new start-target pairs in the generalization phase the modeling suggests that the participants start to either learn the visuomotor mapping since feedback was available or they potentially recalled the mapping, which could have been at least partially learned early on in training. In Experiment 2, when no sequential planning was required and feedback was withheld, performance in the generalization phase remained worse for the Single group compared to the Multiple group. However, their performance was still greater than chance, leaving open the possibility that participants may have recalled (at least partially) the mapping. As such, it remains unclear if the visuomotor mapping is learned to some degree early on and remains latent, which can be later recalled. To test this idea, in Experiment 3, participants briefly trained in the Multiple condition before being exposed for a long period by the Single condition, followed by a generalization phase where new start-target pairs were introduced. If a visuomotor mapping is trained early on and maintained in memory – even if not being used – then participants would show good performance in the generalization phase. Alternatively, if the sequence is ultimately memorized via a model-free algorithm, it could also cause forgetting of the mapping (model-based algorithm) resulting in poor performance in the generalization phase.

#### Behavioral results

In this experiment, participants first performed 80 trials with multiple targets (Multiple trials) followed by a generalization phase of 20 trials. Then, they performed 1000 trials with a single target (Single trials) followed by a second phase of generalization of 20 trials (see Methods) with our primary interest in their performance between the two generalization phases. Importantly, the generalization phase was identical to Experiment 2 where the new targets were only 1 move away and feedback was withheld to prevent learning. We found that prior to starting each generalization phase, the performance of participants was not statistically different between the Multiple or Single trials (Figure 7B; t(27.9) = 1.68, p = 0.104). Importantly, generalization performance was not significantly different in the Single and Multiple trials during training (Figure 7C; t(29.54) = 0.56, p = 0.57). Similarly, there was no significant change in performance from training to generalization in the Multiple (t(15) = 1.05, p = 0.3) or Single (t(15) = -1.43, p = 0.17) trials of the experiment, neither a significant difference among the change in performance in the two trial phases (Figure 7D; t(29.52) = 1.77, p = 0.08). These results suggest that early exposure to variability can allow people to learn the visuomotor mapping and that they can recall it in the future if novel goals appear.

**Figure 7:**
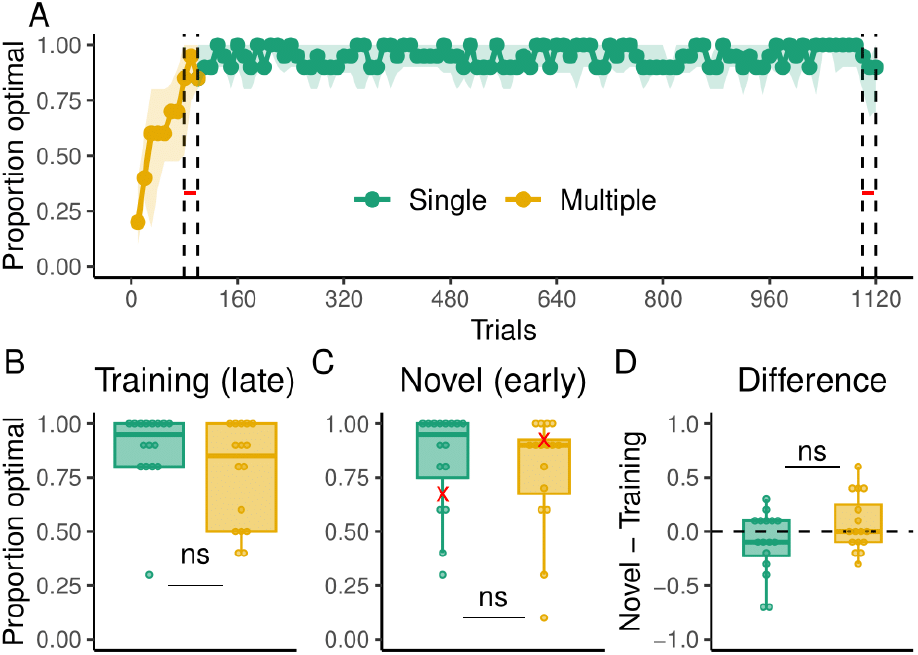
Behavioral results of Experiment 3. **A:** Proportion of optimal arrivals over trials. Participants were exposed to the Multiple trials (gold) followed by the Single group (green). The black dashed line indicates the beginning and end of generalization trials. The red dashed line indicates chance level of performance.The solid dotted line represents the median and the shading the interquartile range. **B:** Proportion of optimal arrivals in the last bin of training trials. **C:** Proportion of optimal arrivals in the first bin of generalization (novel pairs). Red marks indicate performance in the very first trial of generalization for all subjects. **D:** Difference in the proportion of optimal arrivals between the first bin of generalization and the last one of training.

### Experiment 4

It is well-established that skill learning can involve both explicit and implicit processes to varying degrees depending on the training conditions (Jiménez et al., 2006; McDougle et al., 2015; Taylor et al., 2014). In our prior experiments, the visuomotor mapping is relatively simple (i.e., three keys map to three cursor directions) and, as a result, it is possible that participants may have developed explicit knowledge of the visuomotor mapping and/or the sequence of key responses. The latter option is highly likely in the Single conditions in Experiments 1 and 2 where participants only trained to move between one start-target pair. The former is also likely in the Multiple condition where participants could have used an explicit representation of the mapping to simulate action-outcome associations when encountering novel start-target pairs in the generalization phase. To address these possibilities, we introduced a degree of stochasticity in the key-to-direction mapping such that there was a chance on each keypress that the cursor could move to a random direction. Introducing stochasticity between stimulus-response mappings is a common method to blunt awareness and explicit learning in studies of motor sequence learning. By leveraging this method, we ask two questions: 1) Are the performance improvements and generalization observed in the Multiple condition, the result of an explicit or implicit representation of the visuomotor mapping? 2) When explicit learning is blunted, is a visuomotor mapping formed implicitly leading to generalization in the Single condition?

#### Behavioral Results

In this experiment, we aimed to test whether using a stochastic version of the task would blunt explicit learning of the visuomotor mapping, which has proven successful in studies of sequence learning (Jiménez et al., 2006; Schvaneveldt and Gómez, 1998). Two groups of 16 participants performed the grid sailing task as in Experiment 2, in the Single and Multiple conditions, with the exception that the keys moved the cursor according to an underlying key-to-direction mapping with probability 0.8 and to any of the other adjacent directions with probability 0.2. In addition, we evaluated the level of explicit knowledge of the visuomotor mapping at the end of the task by asking participants where they thought the keys move the cursor to. Specifically, pictures of the keyboard keys were displayed on the screen (J, K and L), each of them followed by eight moving options indicated with arrows (top, top-right, right, down-right, down, down-left, left and up-left). Participants had to select among the options the one they believed was the true moving direction of the key (Figure S3).

As a result of the stochasticity in the task, optimal arrivals were rare, therefore as a behavioral measure of performance we considered only arrivals to the target, regardless of the number of key presses. As in Experiments 1 and 2, the Multiple group learned more slowly than the Single group (Figure 8A), but asymptotic performance was similar between the groups by the end of the training phase (Figure 8B; t(26.73) = 1.31, p = 0.19). More importantly, there were no differences in generalization performance between the groups (Figure 8C; t(27.74) = -1.3351, p = 0.1927). While there was no significant difference in the change of performance from training to generalization between the two groups (Figure 8D; t (28.32) = -1.88, p = 0.07), the Single group did significantly decrease its performance from training to generalization phases (t(15) = -2.65, p = 0.01) whereas the Multiple group did not (t(15) = -0.34, p = 0.73).

**Figure 9:**
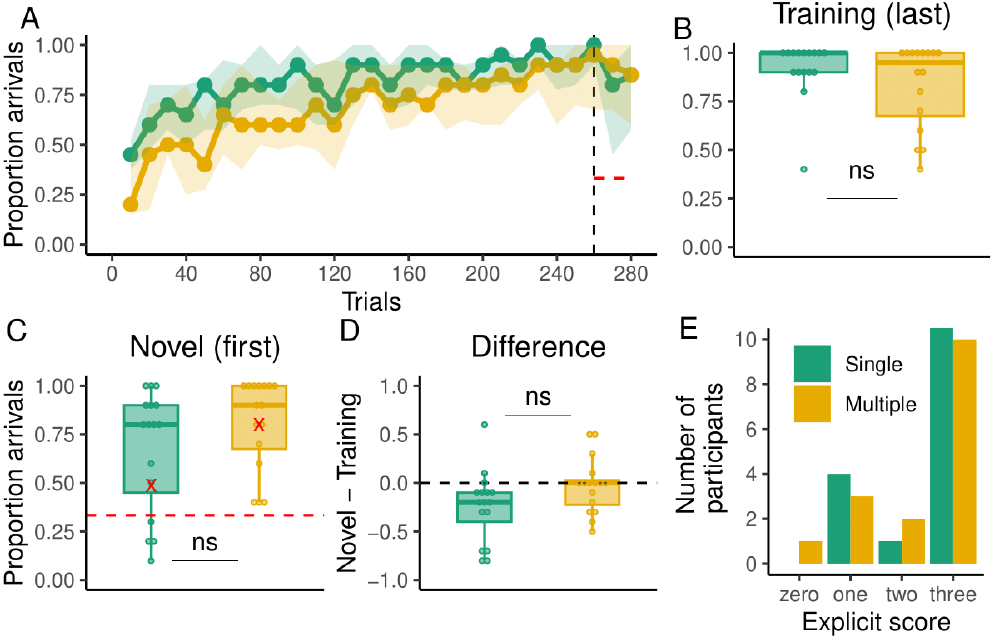
Behavioral results of Experiment 4. **A:** Proportion of arrivals over trials for the Multiple (gold) and Single group (green). The black dashed line indicates the beginning of generalization trials. The red dashed line indicates chance level of performance.The solid dotted line represents the median and the shading the interquartile range. **B:** Proportion of arrivals in the last bin of training trials. **C:** Proportion of arrivals in the first bin of generalization (novel pairs). Red marks indicate performance in the very first trial of generalization for all subjects. **D:** Difference in the proportion of arrivals between the first bin of generalization and the last one of training. **E:** Number of participants that correctly knew zero, one, two or three moving directions of the keys.

At the end of the experiment, participants performed an explicit test where they were asked to report the direction the key moved the cursor to. Both groups showed similar degrees of explicit knowledge of the key-to-direction mapping (Figure 8E). For the Single group 68% (11 out of 16) knew all the keys correctly whereas 32% knew two, one or zero key directions. For the Multiple group 62% (10 out of 16) knew all the keys correctly while 38% (6 out of 16) knew two, one or zero key directions. We further explored whether participants that correctly knew the mapping (scoring 3 in the explicit test) in either group had better generalization performance than people that did not know it, or partially knew it (scoring lower than 3). While participants who had full knowledge of the mapping optimally arrived at the target more often on average than participants who had less knowledge, this difference was not significant (Figure S4). This suggests that explicit knowledge may not be a strong determinant of how people perform in the task.

Overall, the results of Experiment 4 show that both groups had similar explicit knowledge about the mapping and that this knowledge was not related to participants’ generalization performance. In addition, we found that adding stochasticity to the task improves generalization performance in the Single group to the level of the Multiple group, suggesting that the former learned the mapping, potentially by preventing them from memorizing the sequence solution to the goal.

## Discussion

A vast number of skills require the formation of novel visuomotor mappings. Sometimes these mappings can be completely arbitrary like in video games, where an “up” press on a video game controller can lead a virtual character to move or jump. The advantage of learning these mappings, as opposed to simple state-action associations, is that the mappings can be used for planning and generalization to novel contexts (Tolman, 1948; Behrens et al., 2018; Khan, 2018).

Across four experiments, we investigated how training variability can promote the learning of visuomotor mappings in a potentially model task paradigm consisting of moving a cursor to target locations in a grid environment. In Experiments 1 and 2, we found that people who trained with a higher number of start-target pairs (Multiple group) had significantly greater generalization performance as compared to people trained with a single start-target pair (Single group). Critically, in Experiment 2, differences in generalization persisted even when the target locations were only one move away, suggesting that differences in generalization were due to knowledge of the mapping and not due to differences in planning. These results are in line with a large body of research in categorization (Vukatana et al., 2015), language (Lively et al., 1993), problem solving (Pass and Van Merriënboer, 1994) and motor learning (Moxley, 1979), showing that higher variability during training leads to better generalization, although initially making learning more difficult (for a review see Raviv et al., 2022).

We propose that in our experiments this effect is driven in part by the use of different learning processes in the Single and Multiple groups. We predicted that participants exposed to low variability in the Single group would be more likely to rely on a rigid, model-free system. Previous work has shown that in stable environments, this model is a parsimonious solution to learn (Sutton and Barto, 1998). However it fails to adapt to volatile environments (Nassar et al., 2010; Wilson et al., 2013; Ritz et al., 2018; Velázquez et al., 2019) or generalize to novel situations (Daw et al., 2005; Daw et al. 2011). On the other hand, when participants were exposed to higher variability in our Multiple group, we predicted that they would be more likely to learn the underlying mapping of the task, which we represented by a model-based algorithm. This model, in contrast, would be able to generalize well to novel target locations.

Overall, our computational analysis indicates that, when generalization trials were isolated from learning in Experiment 2, most participants in the Single group were best described by a model-free algorithm, whereas most participants in the Multiple group were best described by a model-based algorithm. However, it is relevant to note that a hybrid model also explained an important proportion of our participants’ data, both in Experiment 1 and 2. This is also consistent with previous findings showing that both model-free and model-based systems underlie performance in decision-making and motor learning tasks (Huang et al., 2011; Haith and Krakauer, 2013; Daw et al., 2005; Daw et al., 2011). In situations where both systems can be involved, it is possible to evaluate the relative contribution of each of them to the overall output of the model. When we performed this analysis for Experiment 1, we found that the weight toward the model-based component was significantly higher in the Multiple group for the majority of the training trials and early in generalization (see Figure 4), suggesting that this component had a greater influence in subjects’ responses when variability was higher.

In addition to differences in task performance and model selection, we found that RTs also differed significantly between the Single and Multiple groups in Experiment 1 and 2. In particular, RTs were higher in the Multiple group in the training phase. Previous results have shown that higher RTs could reflect the increased use of model-based, algorithmic computations (Shepard and Metzler, 1971; McDougle and Taylor, 2019). However, differences could also be attributed to people simply having more targets to choose from in the Multiple group as described by Hick’s law (Hick, 1952). Therefore, RT results should not be interpreted in isolation of generalization performance, especially in Experiment 2, and the computational modeling findings.

In Experiment 3, we found that the benefit of having variable training over generalization remained even after a long exposure to no variability. We believe this is a result of the formation of the novel action mapping at an early stage of learning (Fitts and Posner, 1967), and the abrupt change in variability could have separated the mapping memory from future updates (Gershman et al., 2010; Heald et al., 2021), preventing it from being forgotten. Subsequently, during the period of no variability, novel state-action associations could have been formed. If separate memories for the action mapping and the state-action associations were formed, the latter could have potentially been evoked by reducing the preparation time during generalization (Hardwick et al., 2019). Moreover, these results corroborate previous findings that indicate that the benefits of variable training occur when variability is introduced early in learning as opposed to later (Raviv, 2022), but only when it is not too high. In our experiments variable training implied being exposed to four pairs of start-target locations, which were repeated at least 20 times each (early in learning Experiment 3) and up to 70 times (Multiple conditions in Experiments 1 and 2), which we believe provided participants enough familiarization with each of them. Had they experienced more variability, for example by changing the start-target pairs every trial, performance would have been slower and the benefits in generalization could have arrived later.

Finally, in Experiment 4 we discovered that adding stochasticity to the action mapping, prevented the drop in generalization performance observed in Experiments 1 and 2 in the Single group. We believe that the stochastic experimental design prevented participants from completely relying on an explicit memorization of the sequence of keypresses to arrive at the target. Instead, participants likely had to replan on the fly when the cursor moved in an unexpected direction, essentially having the same effect as people that were trained in the Multiple group in Experiments 1 and 2. Importantly, we also found that making the action mapping stochastic had mixed effects on the level of awareness of participants about the true key directions. As shown in Figure S4, the majority of subjects in the Single (68 %) and Multiple group (62%) were able to indicate the actual movement direction of the keys. This contrasts with previous studies on sequence learning where the same level of stochasticity prevented participants from being aware of the underlying sequence. The degree to which this explicit knowledge is necessary for training the mapping needs further investigation.

Overall, our results provide further evidence that people not only can build models about their environment (Tolman, 1948) or their knowledge (Behrens et al., 2018; Constantinescu et al., 2016), but also about their actions (Shadmeher, 2004). Importantly, we have found that this can happen for skills that are built *de novo*. We propose that the acquisition of these models, in the form of action mappings, can be induced by training variability. Crucially, there is currently not an overarching explanation why higher variability typically leads to better generalization, and alternative hypotheses to the one of internal models would have to be ruled out (Raviv, 2022) in future work.

Whereas most of previous studies in sequence learning like SRT, *m* x *n* tasks or discrete sequence production have allowed the study of externally generated sequences specified by the experimenter, there has been a recent interest in sequences that humans generate internally (Fermin et al. 2010, 2016; Bera et al., 2021; Dundon et al., 2022), which, by not being constrained, allow us to explore the planning processes that make humans arrive at given solutions to achieve goals. A model task in this direction has been grid navigation. Our work provides a step in this direction by further providing cognitive models of the processes that might generate these sequences: model-based mapping learning or state-action associations. We believe these types of tasks are good models of a variety of the activities that humans perform in their lives such as playing video-games, musical instruments or sports, where improvisation and self selection of actions is a common feature.

In addition, grid navigation as in the current experiments sits at the intersection of motor learning and spatial navigation where the interaction of procedural and declarative processes likely occurs. Previous studies have highlighted that participants with some impairment in declarative knowledge do not perform as well as controls in similar tasks (Nissen and Bullemer, 1987). Therefore, grid navigation could be used as a testbed for how declarative knowledge contributes to the acquisition of a motor skill. At the same time, it rests at the level of complexity where it is still tractable to build relatively simple cognitive models to explain human performance.

### Limitations

We believe that in order to fully explore the scope of motor skill acquisition, a more complex mapping could be used for future experiments, e.g., one with more actions or a less intuitive movement rule. This would make the learning process more closely resemble the one that people go through in their lives where ceiling performance might be acquired over hours or days of practice, unlike in our experiment where that level was acquired in minutes.

Regarding the cognitive models we explored in Experiment 1 and 2, it can be observed in the absolute goodness of fit that there is still room for improvement at explaining our data sets. As such, other components can be incorporated into our models that would potentially improve the fit. For example, a persistence or switching component that captures responses that do not depend on reward history but on choice history (Daw, 2011; Miller et al., 2019). A key might be more or less likely to be pressed if it was pressed in the past regardless of the outcomes. This autocorrelation among responses can occur within a state (what was pressed the last time the ship appeared in state *s* of the grid?) or across states (what was pressed in the previous moves while navigating across the states of the grid?).

On the other hand, the hybrid model that we tested has a time series of free parameters (weights) that combines the MB and MF components, which makes its complexity grow with the number of trials. Here, our goal was to explore the behavior of the weights with no dependencies between them to let any trend show up on its own, however, alternative reparametrizations are possible were ω_*t*_ is a function of ω_*t*−1_, and the parameters of that function are estimated. This would reduce the complexity of the model and could be explored in future work.

Finally, we have used Breadth First Search as a planning component in our model-based algorithm given its simple implementation and uninformed structure. However, future work could explore the use of path search algorithms that more closely resemble the planning processes of humans. For example, the use of Heuristic-based algorithms would specify that some trajectories towards the goal are more likely to be explored than others using a value function.

This type of search algorithms have been used to explore how humans plan in board games (van Opheusden and Ma, 2019), and their use in grid navigation could improve the models’ fit.

## Methods

### Participants

112 undergraduate students (49 males, 58 females, 4 non-binary and 1 preferred not to say; mean age = 19.9, sd = 1.4) from Princeton University were recruited through the Psychology Subject Pool. Sample sizes were based on prior studies of the grid sailing task (Fermin et al, 2010, 2016; Bera et al., 2021). The experiments were approved by the Institutional Review Board (IRB) and all participants provided informed consent before performing the experiment.

### Apparatus and task design

All experiments were performed in person using the same computer equipment. Stimuli were displayed on a 60 Hz Dell monitor and computed by a Dell OptiPlex 7050’a machine (Dell, Round Rock, Texas) running Windows 10 (Microsoft Co., Redmond, Washington). Participants made their responses using a standard desktop keyboard. All experiments were programmed in CSS, Javascript and HTML, and run on a web browser and hosted on Google Firebase. Subjects were seated in front of the computer and were asked to follow the instructions to begin the task.

We employed a variant of the grid-sailing task based on Fermin et al. (2010) in which participants were required to navigate a cursor from a starting position to a target location on a 9×9 grid using the J, K and L keys of their keyboard. On Experiment 1-3, each key moved the ship deterministically to one of three possible directions: right, down-left or up-left. On Experiment 4, the keys’ directions followed a stochastic rule (see below). At the onset of the experiment, participants were provided with the following instructions *“In this game, you will use the letters J, K and L of your keyboard to move a vehicle through a grid to a target location. Your goal is to arrive using the shortest route. If you arrive with the shortest route, you will see a happy face. If you arrive using a different route, you will see a neutral face. If you do not arrive after a certain time, you will see a sad face*.*”* After participants confirmed they understood the instructions, the task began. The cursor was displayed as a ship and the targets as anchors. Additionally, to make the task more engaging, it was performed with a background of the ocean with quiet wave sounds and ‘bubble’ sounds every time the cursor moved.

On a given trial, the cursor and a target appeared in locations that varied across experiments (see figure 1). Depending on the performance of the trial, subjects could receive three types of feedback. If they did not arrive at the target in less than 10s, a sad face appeared in place of the target, along with a “wrong sound” indicating they had failed. If participants arrived at the target but not in the minimum number of key presses, a neutral face and sound were presented. If they arrived using the minimum number of key presses, a happy face with a “correct sound” was presented. The visual feedback remained on the screen for 1s after which an inter trial interval of 500 ms occurred. Then, the next trial began. The experiment was divided into a training and a generalization phase, which will be described in detail for each experiment below. During the training phase of all experiments, the targets were placed seven moves away from the start location.

### Experiment 1 Procedure

The goal of this experiment was to determine whether specific training regimes promoted the formation of local state-action associations or a visuomotor mapping, which would manifest in a generalization phase. For all participants, the J, K and L keys moved the ship to the down-left, right and up-left, respectively. The training phase consisted of 260 trials which were followed by 20 generalization trials interleaved with 20 training trials, giving a total of 300 trials. Subjects were randomly assigned to one of two groups that differed in the number of start-target pair locations presented during training. In the Single group (n=16), a unique start-target pair was presented for all training trials. In this case, the target could be reached using a unique sequence of key presses (e.g., J-L-J-L-J-L-K), however, participants were not constrained or encouraged to do so.

For the Multiple group (n=16), four start-target pairs were presented throughout training, where each of them appeared 65 times. We randomized the pairs such that the same pair did not show up more than twice in a row and all four pairs appeared once before observing them again. Additionally, the target for each pair could not be reached using the same sequence of key presses that arrived at other targets. In the generalization trials, four novel start-target pairs were presented for both groups. The target was placed seven moves away from the starting point just as in the training trials. Each of the generalization pairs was repeated five times but no pair appeared more than twice in a row, and all four pairs were observed before showing them again. No performance feedback (faces and sounds) was provided in generalization trials.

### Experiment 2 Procedure

In Experiment 2, we tested whether differences in generalization between the Single and Multiple groups in Experiment 1 when error-based feedback was withheld and no (sequential) planning was required. To accomplish this, the target locations during the generalization phase were shown only one step away from the starting point as opposed to seven in Experiment 1. Twelve novel start-target pairs were created by linking four start locations with three possible target locations. All pairs were presented at least once, the remaining eight generalization trials were randomly chosen without replacement from the available twelve. Importantly, in this experiment there were no training trials interleaved with generalization ones. The reason to remove them was to prevent subjects from learning about the mapping with the feedback from the interleaved training trials. This changed the number of trials from 300 in Experiment 1 to 280 in Experiment 2. Additionally, In order to control for mapping-specific effects, in Experiment 2 we randomized the directions each key was assigned to across subjects. The training phase was the same as in Experiment 1 for the Single (n=16) and Multiple (n=16) groups.

### Experiment 3 Procedure

In Experiment 3, we tested whether a short exposure to the training variability followed by a long exposure to no variability would be sufficient to learn the mapping as well as maintain a memory of the mapping, thus affording good performance in the generalization phase at the end of the experiment. Participants (n=16) first trained during 80 trials with four start-target pairs (Multiple trials), then experienced 20 generalization trials with target locations being one move away as in Experiment 2. We chose 80 training trials of training given that in Experiment 1 and 2, asymptotic performance was reached in this time frame by the Multiple group. Following the training phase, participants were exposed to one thousand trials of a single start-target pair (i.e., Single trials), which was chosen randomly from one of the four pairs experienced in the first 80 trials. Finally, they were exposed to a second generalization phase of 20 trials.

### Experiment 4 Procedure

The goal of Experiment 4 was to twofold. First, we sought to test whether the visuomotor mapping was represented explicitly or implicitly. Second, we tested whether training variability induced by stochasticity in the key-to-direction mapping during training would prevent explicit memorization of the sequence and pressure learning of the mapping, which would afford generalization. To accomplish this, we imposed a probabilistic rule over the movement of the cursor. Specifically, during the training phase of both the Single (n=16) and Multiple (n=16) groups, there was a 0.2 probability that the key moved the cursor to any of the other seven directions different from the original mapping. We use this probability based on previous studies on sequence learning that have found that adding this level of stochasticity prevents participants from learning sequences explicitly. In order to evaluate subjects’ awareness of the action mapping of the task, we asked them at the end of the experiment to indicate the direction each key moved the cursor to (Figure S3). Generalization trials were the same as in Experiment 2, with targets being one step away from the start locations.

### Behavioral data analysis

All analyses were performed using the R statistical software (R Core team, 2023) or Matlab version 2022a (Natick, MA: The Math Works, Inc., 2022). Our main behavioral measure was optimal arrivals to the target in Experiments 1-3, which was defined as the minimum number of key presses to move the cursor to the target (7 moves). In Experiment 4, due to stochasticity, our primary measure was simply arrival to the target even if it was not in the minimum number of key presses. We also examined both reaction time (RT), defined as the time between target presentation and the first key press, and inter-keypress interval, defined as the time between each successive key press. These behavioral metrics were binned every ten trials. When relevant comparisons were done between our Single and Multiple groups, we used Welch’s t tests for unequal variances (Welch, 1947).

### Computational modelling

In order to gain mechanistic insight into the learning processes that could have given rise to the results of our experiments, we evaluated three computational models that were fitted to data from Experiment 1 and 2. At one end of the modeling spectrum, we implemented a prediction error RL model to characterize inflexible, habitual behavior, which we believe could be induced in our Single group (model-free). Although this model works in a relatively straightforward manner, it predicts poor generalization as it can only know what to do in situations it has experienced in the past. At the other end of the modeling spectrum, we used a Bayesian model along with a one-step planning process, to represent a learner that acquires the true key-to-direction mapping and leverages it to decide the best course of action (model-based). As we will describe below, this model would be able to generalize well in our task and we believe a similar mechanism could be drawn on in our Multiple group.

#### Model-free (MF)

This model uses prediction errors to update the value of the keys on each grid state. Model-free algorithms have received considerable attention in the past years due to its simple trial-and-error mechanism, which can capture a wide variety of behavioral and neural data (Schultz et al., 1997; Sutton & Barto, 1998; Miller et al., 1995). In our task, it updates the value v for pressing key k after a prediction error δ is observed. More explicitly, for every time step t:

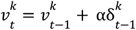

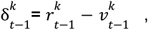

where r is the reward obtained and α is a free parameter that modulates the speed of learning. It is important to note that v is computed for every state and target on the grid, but we have removed those indexes for clarity. We define reward r in terms of the reduction of the chessboard distance d to the target. Specifically:

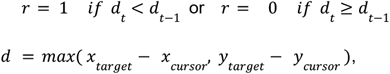

x and y are the grid coordinates of the target and cursor. Then, the probability for pressing key k at time step t is generated using a Softmax function:

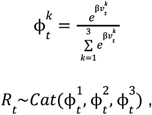

where β is the inverse temperature parameter and R_t_ is the key press at time step t. This model has two free parameters: α and β, for which we specified per-subject prior distributions as α∼Uniform(0,1) and β∼Uniform(0,10). Note that whereas many model-free approaches (temporal-difference methods, etc.) to multi-step decision tasks of this sort recursively learn a multi-step value function measuring distance to goal, here we streamline this approach slightly by defining the target value at each step non-recursively, in terms of the simple chessboard heuristic at each step. This is similar to advantage learning (itself a variant of the actor-critic), but with the value function component fixed as the chessboard distance. We believe that the reduction in the chessboard distance is an intuitive measure of reward in this model, as it is equivalent to visually getting closer to the target. However, this form of distance assumes the cursor can move to any of the adjacent locations, which is not true in our experiments, but is reasonable in an agent that has no knowledge of the key-outcome mapping. As we will see in our next model, the distance to the target can instead be measured as the number of key presses away from it. When the available moves of the cursor are constrained, the key-press distance can differ from the chessboard distance. More importantly, knowing the key-press distance implies knowledge of the true key-outcome mapping, a fundamental property of our next model.

#### Model-based (MB)

In this model, a probability distribution over the key-outcome mapping is updated using Bayes rule and subsequently used to reduce the number of key presses away from the target. In particular, for every key, the cursor movement direction x is assumed to be generated by a Categorical distribution:

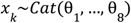

where (θ_1_, …, θ_8_) are the true probabilities that the key k moves the cursor to each of the eight adjacent locations. These probabilities are unknown but can be inferred using Bayes rule. In order to do that, a prior distribution over (θ_1_, …, θ_8_) has to be specified which represents the initial knowledge of the key-outcome mapping. For reasons of conjugacy, it is convenient to choose a Dirichlet distribution:

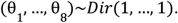

Making the initial parameters equal to 1 gives no preference for any direction *a priori*. Then, the posterior belief about the mapping is described by another Dirichlet distribution:

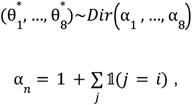

where 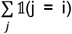 is the number of times the key was observed to go in the ith direction. The expected value of the parameters 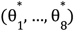 can be computed to have a vector of probabilities π instead of a vector of random variables:

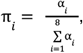

π_*i*_ is the probability that the cursor goes to the ith direction. That is, if a key is pressed, the cursor can end up in the eight adjacent locations with probabilities π. In model-based reinforcement learning π corresponds to the transition probabilities for a given state and action. Our model is a special case of these algorithms for which the transition probabilities are the same for all states. These probabilities are then used to compute the expected distance to the target in the next time step if that key was pressed:

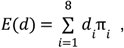

where d is the actual distance to the target, that is, the number of key presses away from it. In order to compute d, we used Breadth First Search (BFS; Erickson, 2019). BFS transforms our grid environment into a graph where each node represents a grid state and nodes are connected among themselves according to the possible transitions in the grid given the action mapping. BFS is thought to represent the planning process in the model-based algorithm and can be implemented in the pseudocode of Algorithm 1.

##### Algorithm 1: BFS algorithm to compute the distance to the target. *Note:* The function *Inside* returns whether the new location is inside the board

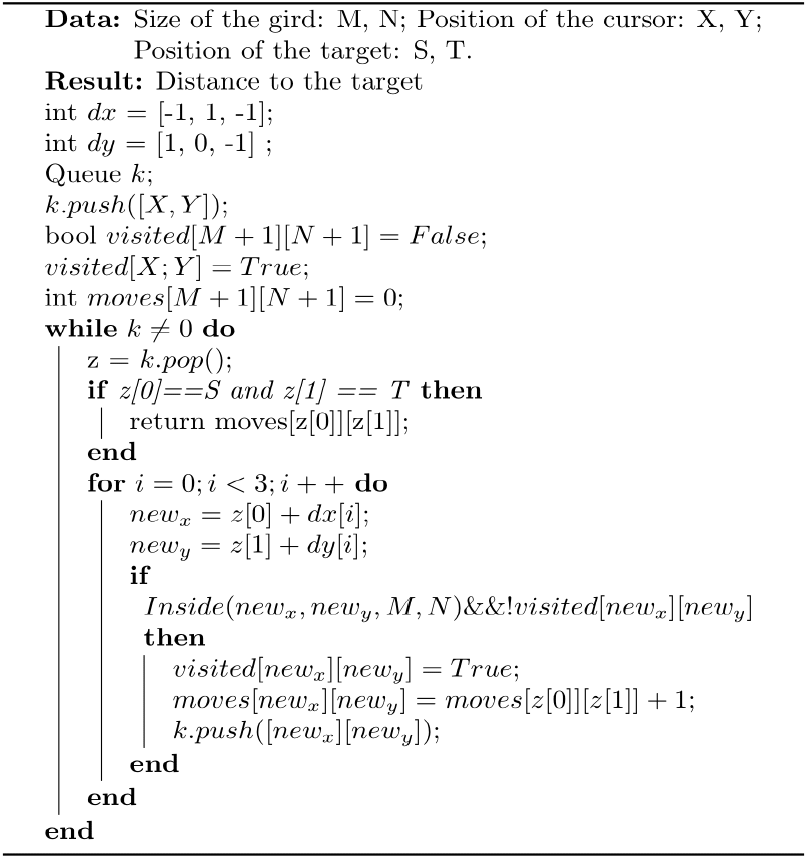

What BFS does is to search on the graph created with the grid environment by first visiting the nodes that are one move away from the current location, then it checks if the target is there; if it isn’t, then it continues searching in the nodes that are two moves away and so on. It continues this process until it reaches the target. We can use − *E*(*d*) to represent the value of pressing a given key. Changing the sign to negative makes lower distances more valuable, then these quantities can be plugged into a Softmax function:

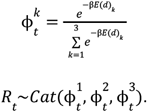

This model has one free parameter: β, for which we set a prior β∼Uniform(0,10).

#### Hybrid model

We considered a third model which is a weighted combination of the RL and Bayesian models:

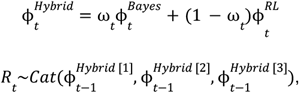

where ω_*t*_ is the weight for the Bayesian component at time *t*. This model has as free parameters: α, β and one weight parameter ω per trial, whose prior distributions are α∼Uniform(0,1), β∼Uniform(0,10) and ω_*t*_ ∼Uniform(0,1).

#### Model evaluation

We approximated the per-subject posterior distributions of parameters of our models using the package JAGS (Just Another Gibbs Sampler; Plummer, 2003) implemented in R code. JAGS uses Markov Chain Monte Carlo to obtain dependent samples from the posterior distribution. For our three models we used four independent chains with 500 samples each, and a burn-in period of 300 samples (samples that were discarded as they likely did not reflect the true distribution). A thinning of 2 was used (i.e., values were taken every 2 samples of the chain) to reduce autocorrelation within chains. Convergence was assessed using the standard potential scale reduction factor 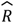 (Gelman and Rubin, 1992) and by visual inspection.

#### Logarithm of the marginal likelihood (LML)

As a metric for model comparison we used the LML. The marginal likelihood (ML), also known as Bayesian evidence, corresponds to the denominator in Bayes rule. It is a measure of how a model explains the current data set and can also be used to compute other model comparison metrics like Bayes factors. For a given model *M*_*i*_, it is described by:

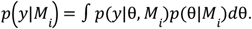

This integral is usually intractable and has to be approximated numerically. To do so, we used a method known as bridge sampling (Gronau et al., 2017a, 2017b) which was implemented by the R package ‘bridgesampling’ and the posterior samples obtained from JAGS (for details, see Gronau et al., 2017a, 2017b).

#### Absolute goodness of fit

In addition to computing the LML, which allows us to compare models among themselves, we wanted to see how well they described the data in the absolute sense, that is, compared to a theoretical (near) upper boundary for any probabilistic model, at least given particular assumptions about exchangeability. This approximate upper limit is represented by the negative entropy (Shannon, 1948; Shen & Ma 2016; Grassberger, 1988, 2003) and is given by:

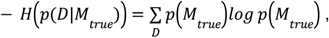

where *p*(*D*|*M*_*true*_) represents the probability distribution of the data given the true model. The negative entropy is a non-positive quantity and intuitively represents how much we can know about the data from the true generative model. An estimator of the negative entropy that has small error even with few data points is given by Grassberger (1988, 2003; see also Shen & Ma, 2016). For our experiments, this estimator is given by:

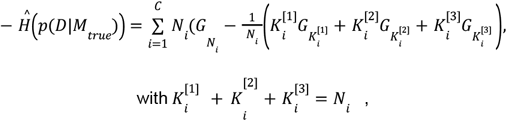

*G*_1_, *G*_2_, … are defined by:

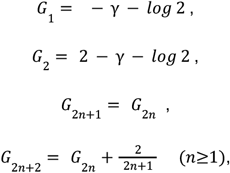

Thus,

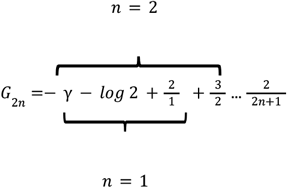

*C* represents partitions of the data, e.g., experimental conditions, which in our case equals the number of unique pairs of start and target locations times the number of states in the grid. Therefore, *C* was not the same in all the experimental groups. In Experiment 1, the number of states in the grid was 81. The number of start and target pairs was five for the Single group and eight for the Multiple group. Therefore, *C* = 405 and *C* = 648 for the Single and Multiple groups, respectively. In Experiment 2, the number of grid states was 81 for both groups. The number of unique pairs of start and target locations was thirteen for the Single group and sixteen for the Multiple group. Therefore, *C* = 1053 in the Single group and *C* = 1296 in the Multiple group. N_i_ is the total number of responses in the partition *i* of the data. K_i_^[1]^ is the number of responses to key 1 in the *i* partition of the data, K_i_^[2]^ the number of responses to key 2 in the *i* partition of the data and K_i_^[3]^ the number of responses to key 3 in the *i* partition of the data. Importantly, this estimator assumes that the distribution of the data given the true model is stationary, which is not necessarily the case of our task as participants’ responses can change due to learning. However, given that subjects’ performance stabilized relatively quickly as we can see in Figure 2 and 5, we considered it would be a reasonable approximation to an upper boundary of the models’ performance.

The negative entropy can be compared with the negative cross-entropy, which intuitively represents how much we can know about the data from an imperfect model (our models). The negative cross entropy is given by:

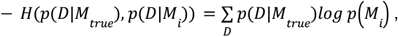

where *p*(*D*|*M*_*i*_) represents the probability distribution of the data given the proposed model *M*_*i*_. The negative cross-entropy is also a non-positive value. An estimator of the negative cross entropy is the logarithm of the likelihood function evaluated at the maximum likelihood estimates of the parameters (Shen & Ma, 2016). In order to obtain this quantity, we used the set of posterior samples that provided the maximum value of this function. In order to provide a simple visualization of the absolute goodness of fit, we computed the proportion of the explainable variability in the data that was explained by the models:

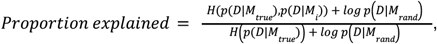

*log p* (*D*|*M*_*rand*_) is the logarithm of the likelihood of the data given a model that assumes all responses are equally likely and represents a lower boundary for all models. In the numerator, we have what *is* explained by a proposed model (as compared to the lower boundary), relative to what *can* be explained (difference between the upper and lower boundary), which is in the denominator.

#### Group model comparison

In addition to comparing the models among themselves at the individual level, we performed group model comparison following Stephen et al. (2009). Following this paper, the probabilities (q_1,_ q_2,_ q_3_) of our models in the population follow a Dirichlet distribution:

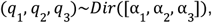

the parameters α = [α_1_, α_2_, α_3_] can be estimated by iterating the following algorithm provided by the authors and that we implemented in R code:

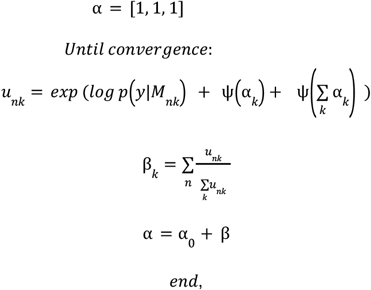

where k is the number of tested models, n the number of subjects and Ψ the digamma function. Importantly, this algorithm only requires that we provide the log marginal likelihoods from each model which we had already computed before for individual model comparison. In order to avoid extremely big numbers from *u*_*nk*_, which returned ∞ in R, we used the logarithm of the marginal likelihood with base 100. We iterated the algorithm 10^3^ times to provide reliable estimates of α. The new parameters of the Dirichlet distribution can be used to compute the probabilities [r_1,_ r_3,_ r_3_] that a randomly selected subject follows any of the tested models:

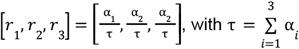

Finally, we computed the probability that a given model k is more likely than the others in the population, i.e., the exceedance probability φ_*k*_:

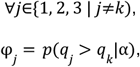

by the Law of Total Probability:

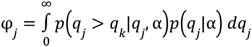

This integral can be approximated numerically by the method provided in Soch and Allefeld (2016) implemented in Matlab code.

#### Parameter recovery

We verified that parameters from our models could be recovered. In order to do so, we generated data from each of them using 100 random samples of their parameter space; then, we performed maximum likelihood estimation using Bayesian Adaptive Direct Search (Acerbi and Ma, 2017) to attempt to recover the parameters generating the data. Finally, we plotted the simulated versus the fitted parameters and computed the Pearson correlation between them. In order to reduce the complexity of the hybrid model, we fitted two weights toward the MB component, one for training trials and one for generalization trials. As can be observed in Figure S5, most of the parameters from the models can be recovered reasonably well, with the exception of the inverse temperature β. One of the reasons this parameter can be difficult to recover is because there might be systematic biases in the data that are considered as noise, e.g., preference for a particular key (Wilson and Collins, 2019). Modeling these biases is one way to improve the recovery of β. We did not perform further reparametrization of the models as we did not have any particular prediction about the behavior of β.

#### Model recovery

In order to verify that the models were identifiable, we simulated 100 data sets from each of the models using random samples of the parameter space. Then fitted those data sets with all the models. In Figure S6 we showed the confusion matrix with the proportion of times that each model was able to recover the data generated by the other models the best according to the Bayesian Information Criterion (Schwarz, 1978). In general, all the models were able to recover their own data better than the other models above 90% of the times.

## Supplementary material

**Figure S1:**
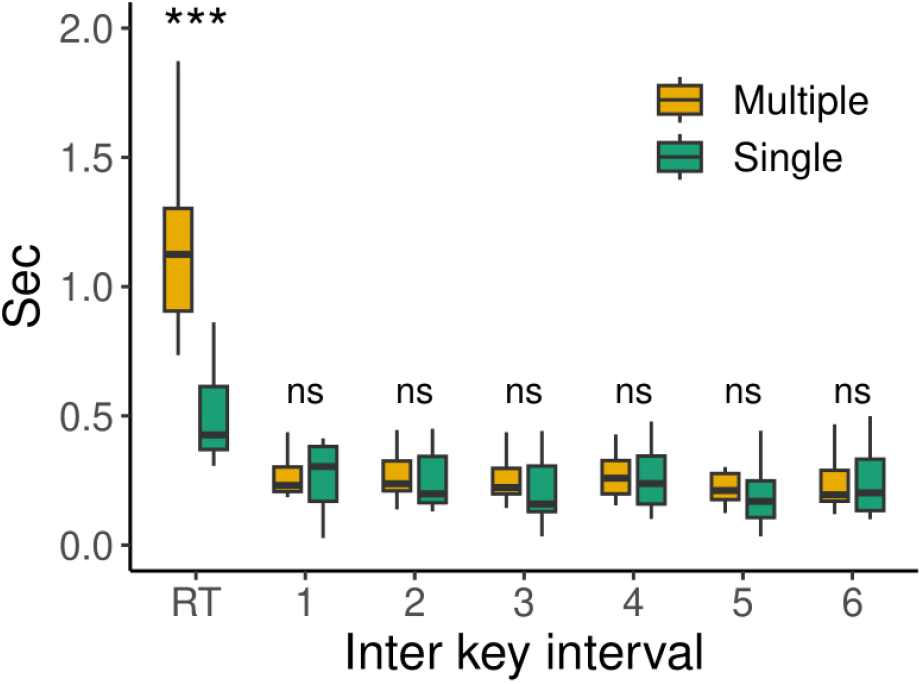
Inter-key-intervals for the first seven key presses.

**Figure S2:**
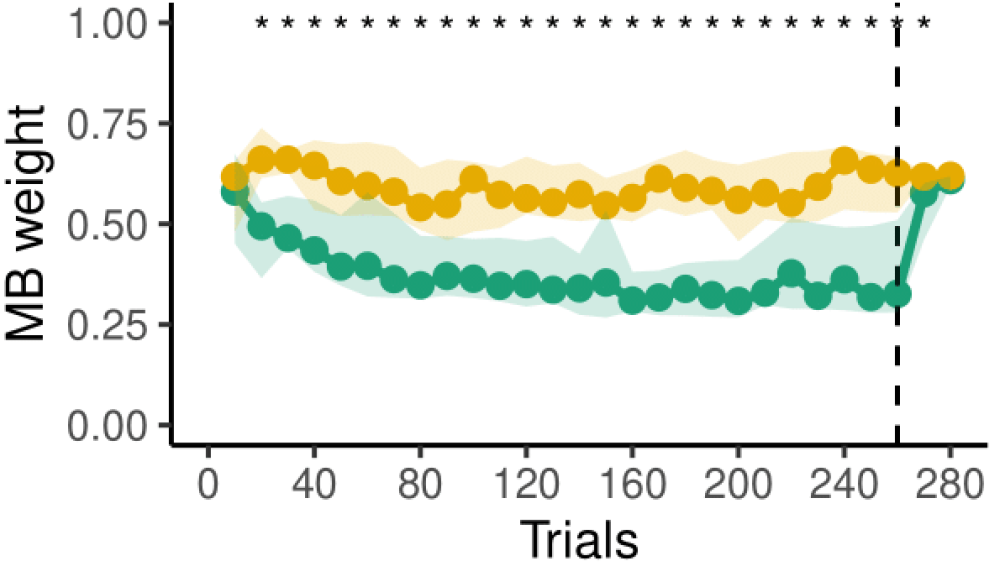
Model-based weight in the Hybrid model for Experiment 2.

**Figure S3:**
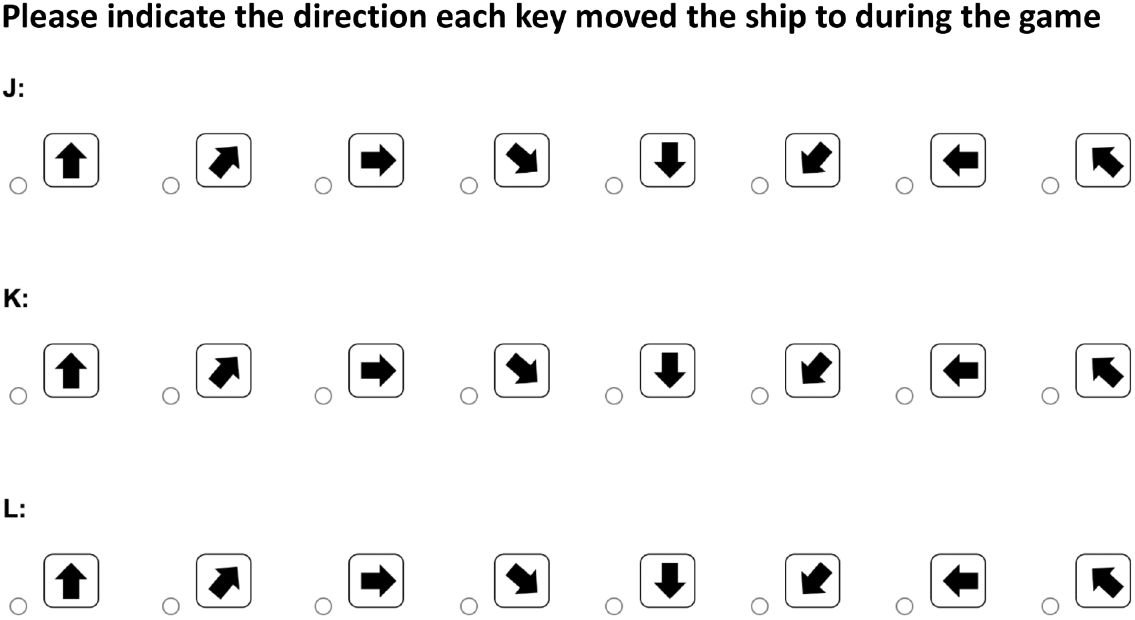
Explicit test of Experiment 4.

**Figure S4:**
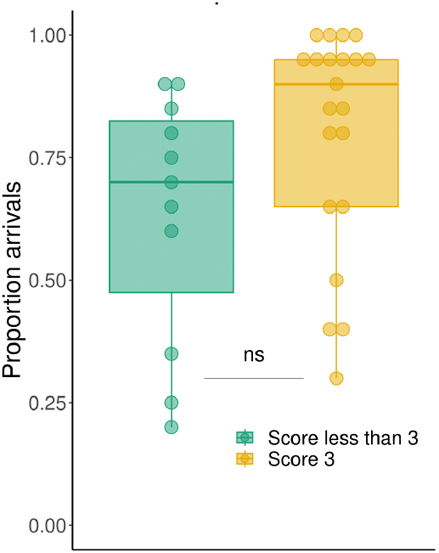
Proportion of arrivals in the generalization phase for participants that score 3 or less than 3 in the explicit test of Experiment 4.

**Figure S5:**
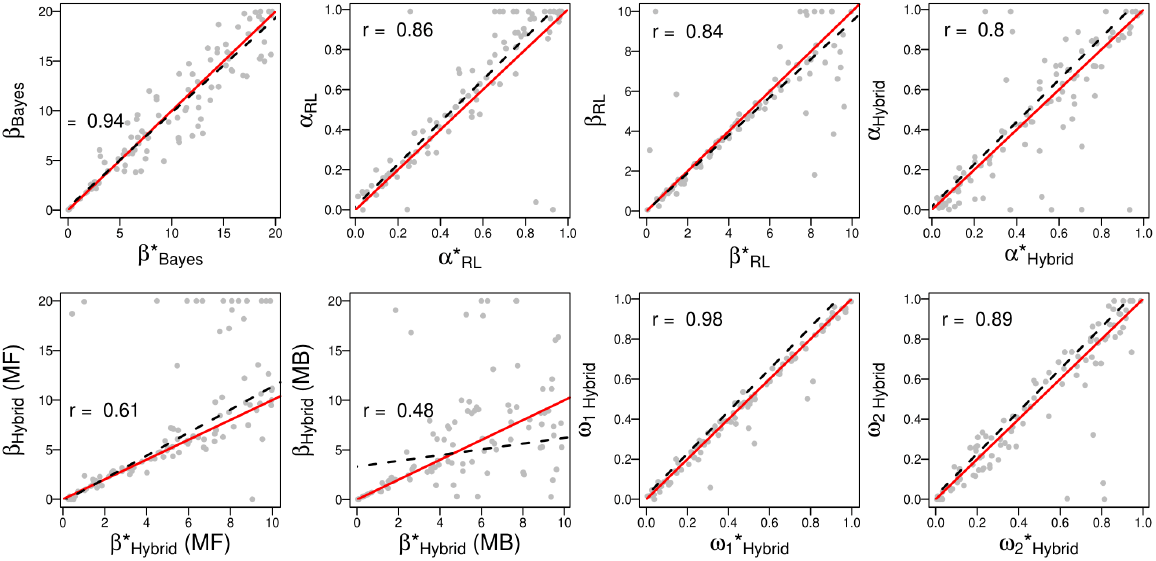
Results of parameter recovery of the Model-free, Model-based and Hybrid models. On the x axis is the simulated parameter and on the y axis the recovered parameter. The subscripts indicate the model. α = learning rate, β =inverse temperature, ω_1_ = model-based weight for training trials, ω_2_ = model-based weight for generalization trials. Red lines represent the identity, the black dashed line is the fit of a linear model to the data and r represents the Pearson correlation.

**Figure S6:**
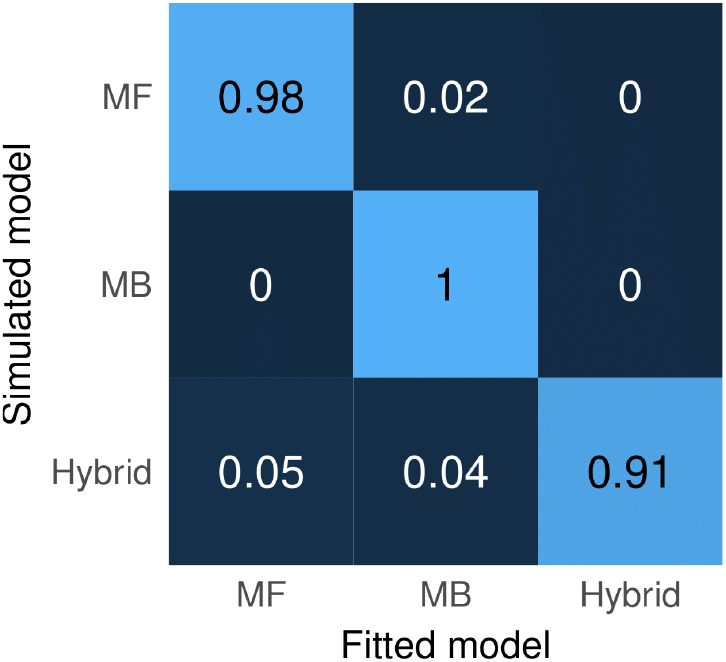
Confusion matrix with the model recovery results. Numbers represent the proportion of times that the model in the Y axis recovered the data generated by the model on the X axis the best according to BIC.

## References

Abrahamse, E. L., Ruitenberg, M. F., De Kleine, E., & Verwey, W. B. (2013). Control of automated behavior: insights from the discrete sequence production task. Frontiers in human neuroscience, 7, 82.

Ackerman, P. L. (1988). Determinants of individual differences during skill acquisition: Cognitive abilities and information processing. Journal of experimental psychology: General, 117(3), 288.

Adams, J. A. (1971). A closed-loop theory of motor learning. Journal of motor behavior, 3(2), 111–150.

Benson, B. L., Anguera, J. A., & Seidler, R. D. (2011). A spatial explicit strategy reduces error but interferes with sensorimotor adaptation. Journal of neurophysiology, 105(6), 2843–2851.

Bapi, R. S., Doya, K., & Harner, A. M. (2000). Evidence for effector independent and dependent representations and their differential time course of acquisition during motor sequence learning. Experimental Brain Research, 132, 149–162.

Bera, K., Shukla, A., & Bapi, R. S. (2021). Cognitive and Motor Learning in Internally-Guided Motor Skills. Frontiers in Psychology, 12, 604323.

Berniker, M., Franklin, D. W., Flanagan, J. R., Wolpert, D. M., & Kording, K. (2014). Motor learning of novel dynamics is not represented in a single global coordinate system: evaluation of mixed coordinate representations and local learning. Journal of neurophysiology, 111(6), 1165–1182.

Behrens, T. E., Muller, T. H., Whittington, J. C., Mark, S., Baram, A. B., Stachenfeld, K. L., & Kurth-Nelson, Z. (2018). What is a cognitive map? Organizing knowledge for flexible behavior. Neuron, 100(2), 490–509.

Catalano, J. F., & Kleiner, B. M. (1984). Distant transfer in coincident timing as a function of variability of practice. Perceptual and motor Skills, 58(3), 851–856.

Constantinescu, A. O., O’Reilly, J. X., & Behrens, T. E. (2016). Organizing conceptual knowledge in humans with a gridlike code. Science, 352(6292), 1464–1468.

Curran, T., & Keele, S. W. (1993). Attentional and nonattentional forms of sequence learning. Journal of Experimental Psychology: Learning, Memory, and Cognition, 19(1), 189.

Daw, N. D., Niv, Y., & Dayan, P. (2005). Uncertainty-based competition between prefrontal and dorsolateral striatal systems for behavioral control. Nature neuroscience, 8(12), 1704–1711.

Daw, N. D., Gershman, S. J., Seymour, B., Dayan, P., & Dolan, R. J. (2011). Model-based influences on humans’ choices and striatal prediction errors. Neuron, 69(6), 1204–1215.

Dundon, N.M., Colas, J.T., Garrett, N., Babenko, V., Rizor, E., Yang, D., MacNamara, M., Petzold, L. & Grafton, S.T., (2022). Decision heuristics in contexts exploiting intrinsic skill. bioRxiv, 2022-04.

Elsner, B., & Hommel, B. (2004). Contiguity and contingency in action-effect learning. Psychological research, 68, 138–154.

Erickson (2019). Algorithms. Independently published.

Estes, W. K., & Burke, C. J. (1953). A theory of stimulus variability in learning. Psychological Review, 60(4), 276.

Fermin, A., Yoshida, T., Ito, M., Yoshimoto, J., & Doya, K. (2010). Evidence for model-based action planning in a sequential finger movement task. Journal of motor behavior, 42(6), 371–379.

Fermin, A. S., Yoshida, T., Yoshimoto, J., Ito, M., Tanaka, S. C., & Doya, K. (2016). Model-based action planning involves cortico-cerebellar and basal ganglia networks. Scientific reports, 6(1), 1–14.

Fitts, P. M., & Posner, M. I. (1967). Human performance.

Gelman, A., & Rubin, D. B. (1992). Inference from iterative simulation using multiple sequences. Statistical science, 457–472.

Gershman, S. J., Blei, D. M., & Niv, Y. (2010). Context, learning, and extinction. Psychological review, 117(1), 197.

Grassberger, P. (1988). Finite sample corrections to entropy and dimension estimates. Physics Letters A, 128(6-7), 369–373.

Grassberger, P. (2003). Entropy estimates from insufficient samplings. arXiv preprint physics/0307138.

Gronau, Q. F., Sarafoglou, A., Matzke, D., Ly, A., Boehm, U., Marsman, M., Leslie, D.S., Forster, J. J., Wagenmakers, E. & Steingroever, H. (2017a). A tutorial on bridge sampling. Journal of mathematical psychology, 81, 80–97.

Gronau, Q. F., Singmann, H., & Wagenmakers, E. J. (2017b). bridgesampling: An R package for estimating normalizing constants. arXiv preprint arXiv:1710.08162.

Hadjiosif, A. M., Krakauer, J. W., & Haith, A. M. (2021). Did we get sensorimotor adaptation wrong? Implicit adaptation as direct policy updating rather than forward-model-based learning. Journal of Neuroscience, 41(12), 2747–2761.

Haith, A. M., & Krakauer, J. W. (2013). Model-based and model-free mechanisms of human motor learning. In Progress in motor control: Neural, computational and dynamic approaches (pp. 1–21). Springer New York.

Hardwick, R. M., Forrence, A. D., Krakauer, J. W., & Haith, A. M. (2019). Time-dependent competition between goal-directed and habitual response preparation. Nature Human Behaviour, 3(12), 1252–1262.

Heald, J. B., Lengyel, M., & Wolpert, D. M. (2021). Contextual inference underlies the learning of sensorimotor repertoires. Nature, 600(7889), 489–493.

Heald, J. B., Lengyel, M., & Wolpert, D. M. (2022). Contextual inference in learning and memory. Trends in Cognitive Sciences.

Hick, W. E. (1952). On the rate of gain of information. Quarterly Journal of experimental psychology, 4(1), 11–26.

Hikosaka, O., Rand, M. K., Miyachi, S., & Miyashita, K. (1995). Learning of sequential movements in the monkey: process of learning and retention of memory. Journal of neurophysiology, 74(4), 1652–1661.

Howard, I. S., Wolpert, D. M., & Franklin, D. W. (2013). The effect of contextual cues on the encoding of motor memories. Journal of neurophysiology, 109(10), 2632–2644.

Howard, I. S., Wolpert, D. M., & Franklin, D. W. (2015). The value of the follow-through derives from motor learning depending on future actions. Current Biology, 25(3), 397–401.

Huang, V. S., Haith, A., Mazzoni, P., & Krakauer, J. W. (2011). Rethinking motor learning and savings in adaptation paradigms: model-free memory for successful actions combines with internal models. Neuron, 70(4), 787–801.

Hunt, L. T., Daw, N. D., Kaanders, P., MacIver, M. A., Mugan, U., Procyk, E., … & Kolling, N. (2021). Formalizing planning and information search in naturalistic decision-making. Nature neuroscience, 24(8), 1051–1064.

Jiménez, L., Vaquero, J. M., & Lupiánez, J. (2006). Qualitative differences between implicit and explicit sequence learning. Journal of experimental psychology: Learning, Memory, and Cognition, 32(3), 475

Jordan, M. I., & Rumelhart, D. E. (1992). Forward models: Supervised learning with a distal teacher. Cognitive science, 16(3), 307–354.

Kerr, R., & Booth, B. (1978). Specific and varied practice of motor skill. Perceptual and motor skills, 46(2), 395–401.

Krakauer, J. W., Hadjiosif, A. M., Xu, J., Wong, A. L., & Haith, A. M. (2019). Motor learning. Compr Physiol, 9(2), 613–663.

Krakauer, J. W., Pine, Z. M., Ghilardi, M. F., & Ghez, C. (2000). Learning of visuomotor transformations for vectorial planning of reaching trajectories. Journal of neuroscience, 20(23), 8916–8924.

Leinen, P., Shea, C. H., & Panzer, S. (2015). The impact of concurrent visual feedback on coding of on-line and pre-planned movement sequences. Acta Psychologica, 155, 92–100.

Liu, X., Mosier, K. M., Mussa-Ivaldi, F. A., Casadio, M., & Scheidt, R. A. (2011). Reorganization of finger coordination patterns during adaptation to rotation and scaling of a newly learned sensorimotor transformation. Journal of neurophysiology, 105(1), 454–473.

Lively, S. E., Logan, J. S., & Pisoni, D. B. (1993). Training Japanese listeners to identify English/r/and/l/. II: The role of phonetic environment and talker variability in learning new perceptual categories. The Journal of the acoustical society of America, 94(3), 1242–1255.

Martin, T. A., Keating, J. G., Goodkin, H. P., Bastian, A. J., & Thach, W. T. (1996). Throwing while looking through prisms: I. Focal olivocerebellar lesions impair adaptation. Brain, 119(4), 1183–1198

Malone, L. A., & Bastian, A. J. (2010). Thinking about walking: effects of conscious correction versus distraction on locomotor adaptation. Journal of neurophysiology, 103(4), 1954–1962.

Martin, T. A., Keating, J. G., Goodkin, H. P., Bastian, A. J., & Thach, W. T. (1996). Throwing while looking through prisms: II. Specificity and storage of multiple gaze—throw calibrations. Brain, 119(4), 1199–1211.

McCracken, H. D., & Stelmach, G. E. (1977). A test of the schema theory of discrete motor learning. Journal of Motor Behavior, 9(3), 193–201.

McDougle, S. D., Bond, K. M., & Taylor, J. A. (2015). Explicit and implicit processes constitute the fast and slow processes of sensorimotor learning. Journal of Neuroscience, 35(26), 9568–9579.

McDougle, S. D., & Taylor, J. A. (2019). Dissociable cognitive strategies for sensorimotor learning. Nature communications, 10(1), 40.

Miall, R. C., & Wolpert, D. M. (1996). Forward models for physiological motor control. Neural networks, 9(8), 1265–1279.

Miller, K. J., Botvinick, M. M., & Brody, C. D. (2017). Dorsal hippocampus contributes to model-based planning. Nature neuroscience, 20(9), 1269–1276.

Miller, R. R., Barnet, R. C., & Grahame, N. J. (1995). Assessment of the Rescorla-Wagner model. Psychological bulletin, 117(3), 363.

Mosier, K. M., Scheidt, R. A., Acosta, S., & Mussa-Ivaldi, F. A. (2005). Remapping hand movements in a novel geometrical environment. Journal of neurophysiology, 94(6), 4362–4372.

Moxley, S. E. (1979). Schema: The variability of practice hypothesis. Journal of motor behavior, 11(1), 65–70.

Mussa-Ivaldi, F. A. (1999). Modular features of motor control and learning. Current opinion in neurobiology, 9(6), 713–717.

Nassar, M. R., Wilson, R. C., Heasly, B., & Gold, J. I. (2010). An approximately Bayesian delta-rule model explains the dynamics of belief updating in a changing environment. Journal of Neuroscience, 30(37), 12366–12378.

Newell, K. M. (1985). Coordination, control and skill. In Advances in psychology (Vol. 27, pp. 295–317). North-Holland.

Newell, K. M. (1991). Motor skill acquisition. Annual review of psychology, 42(1), 213–237.

Newell, K. M., & Shapiro, D. C. (1976). Variability of practice and transfer of training: Some evidence toward a schema view of motor learning. Journal of Motor Behavior, 8(3), 233–243.

Nissen, M. J., & Bullemer, P. (1987). Attentional requirements of learning: Evidence from performance measures. Cognitive psychology, 19(1), 1–32.

Oh, Y., & Schweighofer, N. (2019). Minimizing precision-weighted sensory prediction errors via memory formation and switching in motor adaptation. Journal of Neuroscience, 39(46), 9237–9250.

Paas, F. G., & Van Merriënboer, J. J. (1994). Variability of worked examples and transfer of geometrical problem-solving skills: A cognitive-load approach. Journal of educational psychology, 86(1), 122.

Petrides, M. (1997). Visuo-motor conditional associative learning after frontal and temporal lesions in the human brain. Neuropsychologia, 35(7), 989–997.

Plummer, M. (2003). JAGS: A program for analysis of Bayesian graphical models using Gibbs sampling. In Proceedings of the 3rd international workshop on distributed statistical computing (Vol. 124, No. 125.10, pp. 1–10).

Raviv, L., Lupyan, G., & Green, S. C. (2022). How variability shapes learning and generalization. Trends in cognitive sciences.

Ritz, H., Nassar, M. R., Frank, M. J., & Shenhav, A. (2018). A control theoretic model of adaptive learning in dynamic environments. Journal of cognitive neuroscience, 30(10), 1405–1421.

RStudio Team (2023). RStudio: Integrated Development for R. RStudio, PBC, Boston, MA URL http://www.rstudio.com/.

Schvaneveldt, R. W., & Gomez, R. L. (1998). Attention and probabilistic sequence learning. Psychological Research, 61(3), 175–190.

Schmidt, R. A. (1975). A schema theory of discrete motor skill learning. Psychological review, 82(4), 225.

Shadmehr, R., & Mussa-Ivaldi, F. A. (1994). Adaptive representation of dynamics during learning of a motor task. Journal of neuroscience, 14(5), 3208–3224.

Shadmehr, R., Smith, M. A., & Krakauer, J. W. (2010). Error correction, sensory prediction, and adaptation in motor control. Annual review of neuroscience, 33, 89–108.

Shannon, C. E. (1948). A mathematical theory of communication. The Bell system technical journal, 27(3), 379–423.

Shen, S., & Ma, W. J. (2016). A detailed comparison of optimality and simplicity in perceptual decision making. Psychological review, 123(4), 452.

Shepard, R. N., & Metzler, J. (1971). Mental rotation of three-dimensional objects. Science, 171(3972), 701–703.

Schultz, W., Dayan, P., & Montague, P. R. (1997). A neural substrate of prediction and reward. Science, 275(5306), 1593–1599.

Schwarz, G. (1978). Estimating the dimension of a model The Annals of Statistics 6 (2), 461–464. URL: http://dx.doi.org/10.1214/aos/1176344136.

Simon, D. A., & Daw, N. D. (2011). Neural correlates of forward planning in a spatial decision task in humans. Journal of Neuroscience, 31(14), 5526–5539.

Soch, J., & Allefeld, C. (2016). Exceedance Probabilities for the Dirichlet Distribution. arXiv preprint arXiv:1611.01439.

Stephan, K. E., Penny, W. D., Daunizeau, J., Moran, R. J., & Friston, K. J. (2009). Bayesian model selection for group studies. Neuroimage, 46(4), 1004–1017

Sutton, R. S., & Barto, A. G. (1998). Introduction to reinforcement learning (Vol. 135, pp. 223–260). Cambridge: MIT press.

Taylor, J. A., Krakauer, J. W., & Ivry, R. B. (2014). Explicit and implicit contributions to learning in a sensorimotor adaptation task. Journal of Neuroscience, 34(8), 3023–3032.

The MathWorks Inc. (2022). MATLAB version: 9.13.0 (R2022b), Natick, Massachusetts: The MathWorks Inc. https://www.mathworks.com.

Thorndike, E. L. (1927). The law of effect. The American journal of psychology, 39(1/4), 212–222.

Tolman, E. C. (1948). Cognitive maps in rats and men. Psychological review, 55(4), 189.

van Opheusden, B., & Ma, W. J. (2019). Tasks for aligning human and machine planning. Current Opinion in Behavioral Sciences, 29, 127–133.

Verwey, W. B. (2001). Concatenating familiar movement sequences: The versatile cognitive processor. Acta psychologica, 106(1-2), 69–95.

Vukatana, E., Graham, S. A., Curtin, S., & Zepeda, M. S. (2015). One is not enough: Multiple exemplars facilitate infants’ generalizations of novel properties. Infancy, 20(5), 548–575.

Welch, B. L. (1947). The generalization of ‘STUDENT’S’problem when several different population varlances are involved. Biometrika, 34(1-2), 28–35.

Willingham, D. B., Nissen, M. J., & Bullemer, P. (1989). On the development of procedural knowledge. Journal of experimental psychology: learning, memory, and cognition, 15(6), 1047.

Wilson, R. C., & Collins, A. G. (2019). Ten simple rules for the computational modeling of behavioral data. Elife, 8, e49547.

Wilson, R. C., Nassar, M. R., & Gold, J. I. (2013). A mixture of delta-rules approximation to bayesian inference in change-point problems. PLoS computational biology, 9(7), e1003150.

Velázquez, C., Villarreal, M., & Bouzas, A. (2019). Velocity estimation in reinforcement learning. Computational Brain & Behavior, 2, 95–108.

Yang, C. S., Cowan, N. J., & Haith, A. M. (2021). De novo learning versus adaptation of continuous control in a manual tracking task. elife, 10, e62578.

